# Activity-dependent alteration of early myelin ensheathment in a developing sensory circuit

**DOI:** 10.1101/2021.04.18.440312

**Authors:** Zahraa Chorghay, David MacFarquhar, Vanessa J. Li, Sarah Aufmkolk, Anne Schohl, Paul W. Wiseman, Ragnhildur Thora Káradóttir, Edward S. Ruthazer

## Abstract

Adaptive myelination has been reported in response to experimental manipulations of neuronal activity, but the links between sensory experience, corresponding neuronal activity, and resultant alterations in myelination require investigation. To study this, we used the *Xenopus laevis* tadpole, which is a classic model for studies of visual system development and function because it is translucent and visually responsive throughout the formation of this retinotectal system. Here, we report the timecourse of early myelin ensheathment in the *Xenopus* retinotectal system using immunohistochemistry of myelin basic protein (MBP) along with third-harmonic generation (THG) microscopy, a label-free structural imaging technique. Characterization of the myelination progression revealed an appropriate developmental window to address the effects of early patterned visual experience on myelin ensheathment. To alter patterned activity, we showed tadpoles stroboscopic stimuli and measured the calcium responses of retinal ganglion cell axon terminals. We identified strobe frequencies that elicited robust versus dampened calcium responses, reared animals in these strobe conditions for 7 d, and subsequently observed differences in the amount of early myelin ensheathment at the optic chiasm. This study provides evidence that it is not just the presence but also to the specific temporal properties of sensory stimuli that are important for myelin plasticity.

## Introduction

The development and function of brain circuits relies crucially upon precise timing of neuronal inputs. By insulating axons to regulate the conduction velocity of these inputs, myelination may optimize temporal control over information processing, with implications for synchrony in vertebrate circuits(1, 2). Effects of experience and training on biomarkers of myelination have been reported with white matter imaging techniques and with cellular level investigations(3, 4). These cellular level changes have been studied using extreme manipulations of axonal activity or vesicular release, including sensory deprivation, chronic pharmacological treatment, genetic manipulations, and electrical and optogenetic stimulation(3-5). However, the links between patterns of sensory experience, corresponding neuronal activity, and myelination have yet to be fully elucidated. To study how sensory patterned activity alters myelination during circuit development, we took advantage of the *Xenopus* retinotectal system, which is amenable to imaging, shows precocious visual responsiveness, and has been extensively studied in the context of the effects of patterned activity on synaptic plasticity, structural remodeling, and topographic circuit refinement(6).

## Results and Discussion

To observe myelin ensheathment, we used an antibody against myelin basic protein (MBP), which showed expected band sizes for MBP isoforms (19 and 22 kDa) on Western blots of adult *Xenopus* brain lysate, in accordance with the reported molecular weights of MBP isoforms in Xenopus(7) and other species(8) (Fig 1A). The pattern of MBP immunostaining reflected the laminar organization of the adult optic tectum(9) (Fig 1B), and in the hindbrain was similar to immunostaining for myelin proteolipid protein reported in stage 49 tadpoles(10) (Fig 1C). We cross-validated immunostaining with third-harmonic generation (THG) microscopy, an emerging label-free technique that has been used to image the presence of myelin in the peripheral and central nervous systems(11, 12). THG microscopy reveals sub-micrometer heterogeneities produced at optical interfaces, allowing it to be used as a structural imaging tool in unstained samples(13). Strong THG signal was observed in a subset of MBP-positive fibers and increased at later developmental stages in both the hindbrain (Fig 1D) and optic chiasm (Fig 1E), consistent with the developmental progression from new ensheathment by MBP-positive processes to increasingly compact myelin, giving stronger THG signal. Because MBP expression was highly specific and preceded the onset of robust THG signal, we used MBP immunostaining for the rest of this study to investigate effects from the onset of myelination.

**Figure 1.**
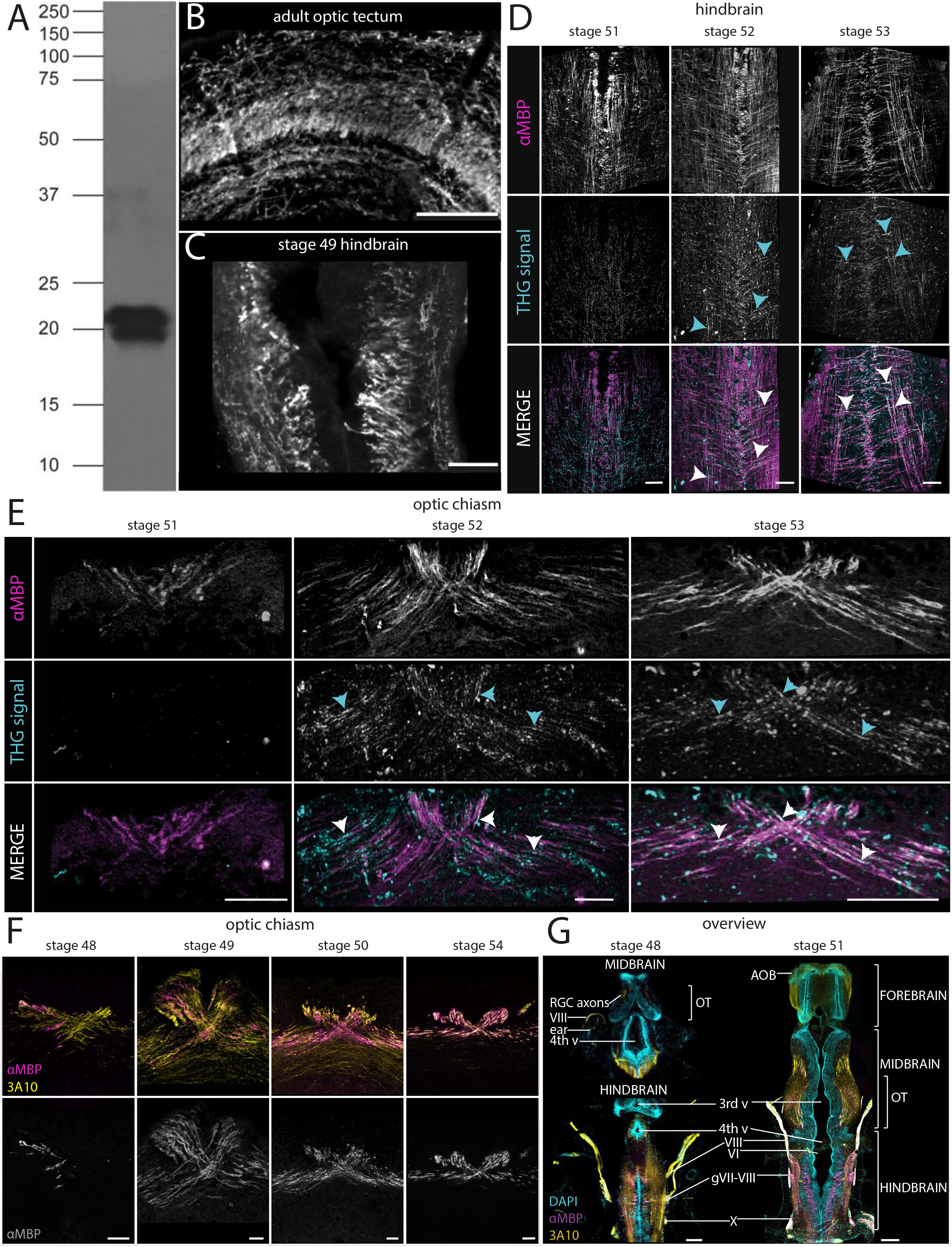
Investigation of early myelin ensheathment in the *Xenopus laevis* tadpole retinotectal system. (A) Western blot using anti-MBP antibody on *Xenopus* adult brain lysate shows expected band sizes of 19 kDa and 22 kDa. (B) THG microscopy of the optic chiasm, showing two-photon MBP immunofluorescence aligned to THG signal in the same sections of tadpole stages 51 to 53. THG signal intensifies later in development than MBP expression, suggesting more compact myelin. Arrows: examples of fibers with both THG and MBP immunofluorescence signal. Colors: αMBP (magenta), THG (cyan). Scale bar: 40 µm. (C, D) Developmental progression of early myelin ensheathment in the retinotectal system, including the optic (C) chiasm beginning at stage 48 and (D) tectum starting at stage 51. Scale bar: 20 µm. (E) The caudo-rostral progression of myelin expression in the brain between stages 48 and 51. Scale bar: 100 µm. (C-E) Colors: αMBP (magenta, grey), 3A10 (yellow), DAPI (cyan). (B-E) Horizontal sections, oriented with rostral toward the top of the panel.

We studied MBP expression in stage 48 to 54 tadpoles, a relevant developmental period when tadpoles have just transitioned from relying on their yolk sack for nutrition to active feeding(14) and show more complex sensorimotor behaviors in response to environmental cues. Immunostaining for MBP alongside monoclonal antibody 3A10 staining of a neurofilament-associated antigen that preferentially labels retinal ganglion cell (RGC) axons(15) and a subset of reticulospinal projections(16) revealed myelination progression in the optic chiasm (Fig 1F). Overall, MBP expression follows a caudal-to-rostral progression in the tadpole brain, highlighted by comparing changes between stage 48 and stage 51 (Fig 1G).

Based on our characterization of myelin ensheathment, we identified stage 48 as the appropriate developmental stage in which to investigate how visual experience modulates MBP expression. Strobe rearing has been previously shown to synchronize RGC firing and modulate topographic map refinement in the optic tectum of Xenopus(17) and in goldfish(18). We used various frequencies of stroboscopic stimuli (“strobe”) to physiologically induce temporally patterned activity in the retinotectal system (Fig 2A). When animals were exposed to strobe, robust calcium responses in RGC axons were evoked by 0.0625 Hz (“slow”) but not by 1 Hz (“fast”) strobe (Fig 2C-F). The Fourier power spectra for these calcium responses showed peaks corresponding to the specific frequencies for slow (Fig 2G) and fast (Fig 2H) strobe. Comparing the power at the stimulus frequencies revealed significantly more power associated with the slow than the fast strobe in each of the animals measured (Fig 2I), most likely due to the phenomenon of temporal frequency adaptation(19).

**Figure 2.**
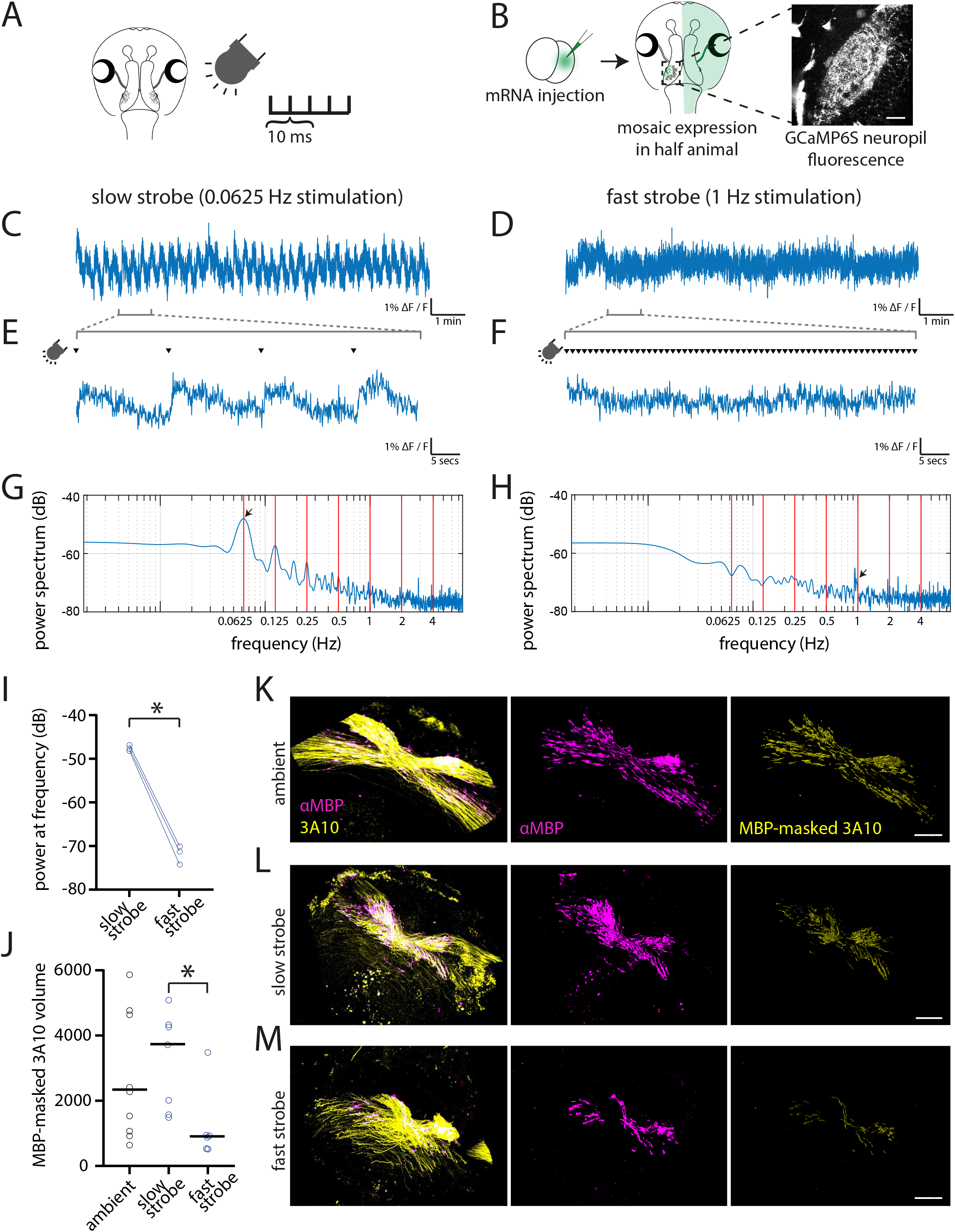
Activity-dependent effects on MBP expression in the retinotectal system with stroboscopic visual stimulation. (A) *Xenopus* tadpoles were exposed to 10 ms stroboscopic flashes at a range of frequencies. (B) For two-photon calcium imaging of RGC axon terminals in the optic tectum, tadpoles with bilateral mosaic GCaMP6s expression restricted to half the animal were generated by microinjection of GCaMP6s mRNA into one blastomere at the two-cell stage of development. Scale bar: 40 µm. (C,D) Representative tectal neuropil axonal calcium responses to the (C) slow strobe (0.0625 Hz) and (D) fast strobe (1 Hz) stimuli over 10 min and zoomed in to a (E,F) 1 min period. Arrowhead: LED flashes during strobe stimulation. (G, H) Fourier power spectra of calcium responses in (C, D). Arrow: peak in power corresponding to strobe frequency. (I) Power at the stimulus frequency was significantly greater for slow rather than fast strobe in each animal (n = 3, paired t-test, *p = 0.0034). (J-M) Animals were raised under ambient light, slow or fast strobe for 7 d, and their optic chiasm digitally reconstructed in 3D. (J) Quantification of the MBP-masked 3A10 volume showed differences between the three conditions. n = 9 for ambient, 7 for slow strobe, 6 for fast strobe. Line: median. Kruskal-Wallis test: *p = 0.0187, Dunn’s test for multiple comparisons: *p = 0.0288 for slow versus fast strobe. (K – M) Representative snapshots of the reconstructions for the (K) ambient, (L) slow, and (M) fast strobe conditions. Colors: αMBP (magenta), DAPI (cyan), 3A10 (yellow). Scale bar: 20 µm.

We therefore raised animals for 7 d under slow or fast strobe conditions or dim ambient light. We chose ambient light as the control since darkness is known to lead to spontaneous local bursting that can drive waves of correlated retinal activity even in the absence of vision(20). Tadpoles were fixed and immunostained, and high-resolution confocal stack images of the optic chiasm were acquired. From these photomontages, 3D digital reconstructions of MBP and 3A10 profiles were used to quantify the total volume of MBP-associated RGC axons at the chiasm (Fig 2J–M). Significantly greater axonal volume was associated with MBP in animals reared under slow versus fast strobe (Fig2J). Overall, our data show that slow strobe, which reliably evoked robust non-adapting calcium responses in RGC axons, increased the axonal volume associated with MBP, whereas fast strobe, which was associated with weak, adapting calcium responses, produced a reduction in ensheathed axonal volume at the chiasm. Dim ambient light-reared tadpoles showed an intermediate level of myelin ensheathment.

Our observation that sensory experience alters myelin is consistent with previous reports(3-5). The differential effect of firing frequency on myelination has been shown with electrical stimulation in co-cultures of dorsal root ganglion neurons and Schwann cells(21) and in the corpus callosum(22). Our findings extend this literature, showing that patterned *sensory experience* of the organism affects MBP expression, with increased myelination under stimulus conditions that elicit elevated, repetitive axonal firing. An ongoing debate is whether activity-dependent myelination may merely reflect changes in axonal morphology, including arbor elaboration(23) or stability of axon terminals(24) rather than neuronal activity itself. However, since we studied myelination along the axonal tract at the optic chiasm, changes in MBP expression here are unlikely to be influenced by axonal terminal branch morphogenesis.

That patterned activity can affect myelin ensheathment hints at the possibility of shared mechanisms for the control of timing-dependent plasticity and myelination during circuit development. Stimulation of spinal cord axons in the zebrafish affects calcium transients in contacting oligodendrocytes, which then predict myelin sheath dynamics(25, 26), reminiscent of correlations between calcium and structural dynamics of neurons for activity-dependent plasticity. Furthermore, specific patterns of neuronal activity induce distinct programs of gene expression(27), conceivably leading to changes in the expression of myelin-related genes. By regulating the speed of action potentials, myelin plasticity may contribute to the precise temporal synchrony and to oscillations in functional circuits(1), as supported by computational modelling: changes in conduction velocity through adaptive myelination allow for more robust synchronization of the network than can be accounted for by synaptic gain alone(2). Lastly, disorders such as autism and schizophrenia have been linked to myelin abnormalities and to dysregulation of temporal synchrony, highlighting the potential importance of experience-dependent myelination in circuit development and refinement.

## Acknowledgements

We thank Drs. Alyson Fournier, Barbara Morquette, Nelly Vuillemin, Nicholas Marsh-Armstrong, and Tim Kennedy for their scientific advice, as well as the MNI Microscopy Core Facility and the McGill Advanced Bioimaging Facility. This work was funded by Fonds de recherche du Québec– Santé (Grant 31036; ER), MNI–Cambridge Collaboration Grant (RTK & ER) and Douglas Avrith Award (ZC), European Research Council Horizon 2020 Research and Innovation Programme (Grant 771411; RTK), Natural Sciences and Engineering Research Council of Canada (NSERC) Discovery Grant to PWW, and NSERC and McGill University Integrated Program in Neuroscience Scholarships to ZC and VJL.

## Materials and Methods

### Experimental Model and Subject Details

All procedures were approved by the Animal Care Committee of the Montreal Neurological Institute at McGill University in accordance with Canadian Council on Animal Care guidelines. For the developmental progression, tadpoles were acquired from Boreal Science (RRID:XEP_Xla100). Upon receiving them, they were staged as per Nieuwkoop and Faber(14) and immediately fixed for sectioning and immunostaining.

For the calcium imaging experiment, we generated bilateral mosaic tadpoles, with fluorescent protein expression restricted to one-half of the animal (described in detail by Benfey et al(28)). Briefly, female albino *Xenopus laevis* frogs (RRID:XEP_Xla300) from our in-house breeding colony were primed with 50 IU pregnant mare serum gonadotropin (PMSG; Prospec Bio HOR-272), and 3 d later, were injected with 400 IU human chorionic gonadotropin (hCG; Sigma-Aldrich CG10; RRID:SCR_018232). The following day, eggs from primed females were collected for in vitro fertilization and co-injection of GCaMP6s and mCherry messenger RNA (mRNA) into one blastomere of two-cell stage embryos. Several days after injection, we screened for animals with unilateral mCherry and high levels of GCaMP fluorescence for use in calcium imaging experiments.

For the strobe-rearing experiments, 2-3 d after priming females with 50 IU PMSG, 400 IU hCG was injected into females and 150 IU into males to induce mating. Fertilized eggs were collected the following day and raised in 0.1X Modified Barth’s Solution with HEPES (MBSH) prepared from 10X MBSH stock solution (88 mM NaCl, 1 mM KCl, 2.4 mM NaHCO3, 0.82 mM MgSO4 x 7H2O, 0.33 mM Ca(NO3)2 x 4H2O, 0.41 mM CaCl2, 10 mM HEPES, pH 7.4). Once animals reached stage 48, they were placed in a 6-well plate in a box that blocked out external light, while exposed to a LED array composed of green luxeon LEDs controlled by a Master 8 Stimulator (A.M.P.I.). Animals were exposed to stroboscopic stimuli with 10 ms full-field light flashes presented at different frequencies or to constant illumination at the same intensity (“ambient”) for 7 d. MBSH media and Sera-Micron 50ml growth food for fish and amphibians (Sera) was refreshed every 1-2 d.

### Western blot

Samples were prepared from adult male Xenopus brain. After adult males were anesthetized in 0.2% MS-222 (Sigma A5040) and decapitated, the brain was dissected and homogenized in RIPA extraction buffer (10mM HEPES/NaOH pH 7.4, 150mM NaCl, 1% Triton X-100, 1% SDS) with protease inhibitors (Calbiochem Protease Inhibitor Set V EDTA-free). Western blot analysis was performed with Bio-Rad wet transfer system and PVDF membrane (Millipore). The rat anti-MBP antibody (Abcam [clone 12] ab7349; RRID:AB_305869) was used at 1:5000 in 5% skim milk, allowing the blot to incubate overnight at 4°C. Incubation with the secondary antibody, goat anti-rat HRP (Jackson Immunoresearch 112-035-167; RRID:AB_2338139), was performed at 1:20 000 in 5% skim milk for 1 h at room temperature. The blots were developed with Immobilon Western chemiluminescent HRP substrate (WBKLS0500).

### Immunohistochemistry

Animals were anesthetized in 0.02% MS-222, fixed by immersion in 4% paraformaldehyde (Cedarlane (EMS) 15735-30-S) in PBS for 1 hr at room temperature, transferred to ice-cold 100% methanol, and post-fixed overnight at –20oC. Samples were then washed for 1 h in a solution of 100 mM Tris/HCl, pH7.4 with 100 mM NaCl. Infiltration and cryoprotection of the samples was performed by incubation overnight at room temperature in a solution of 15% fish gelatin (Norland HP-03) with 15% sucrose, and subsequently repeated in 25% fish gelatin with 15% sucrose. Samples were embedded and frozen in a solution of 20% fish gelatin with 15% sucrose for cryosectioning. Sections were acquired in the horizontal orientation at 20 µm thickness on a cryostat and directly mounted onto Superfrost-plus slides (Fisher).

Slides were incubated with blocking solution (10% bovine serum albumin and 5% normal goat serum in PBS). Sections were immunostained with rat anti-MBP antibody (1:200; Abcam [clone 12] ab7349; RRID:AB_305869), mouse 3A10 (1:400; DSHB Hybridoma Product 3A10; RRID:AB_531874), and cell nuclei counterstained with DAPI (1:1000; Invitrogen D-1306; RRID:AB_2629482). The 3A10 monoclonal antibody is neuron-specific(29), preferentially labelling a subset of hindbrain spinal cord projecting neurons(30) and RGC axons(15). The secondary antibodies used were goat anti-rat IgG Cy3 (1:200; Jackson Immunoresearch 112-165-175; RRID:AB_2338252) and goat anti-mouse IgG Alexa-647 (1:200; Invitrogen A21236; RRID:AB_2535805). We used highly cross-adsorbed secondary antibodies in all instances to prevent cross-reactivity between mouse and rat primary antibodies. Images were acquired with 10x/0.40 CS, 40x/1.30 oil CS2, or 63x/1.40 oil CS2 objectives at a Leica SP8 confocal microscope.

### Third Harmonic Generation Microscopy

Third harmonic generation (THG) imaging was performed on a custom-built laser scanning microscope with forward detection as described in detail by Kazarine et al(31). The setup used a customized upright multiphoton microscope (FV1200 MPE, Olympus Canada Inc, ON, Canada) equipped with a motorized stage and a 25x water immersion objective (1.05 NA; 2 mm working distance; XLPL25XWMP(F), Olympus Canada Inc, ON, Canada). Samples were excited by a Ti:Sapphire laser (Mira 900F, Coherent, CA) pumped by a 532 nm laser (Verdi V18, Coherent, CA). The excitation laser provides 200 fs pulses at a 76 MHz repetition rate and feeds into an optical parametric oscillator (Mira OPO, Coherent, California, U.S.A.), enabling 1150 nm pulses with a femtosecond pulse length necessary to serve the momentum conservation law (phase matching condition) for THG. 50 mW continuous power was measured at the plane of the sample. The emission light was split spectrally by a dichroic mirror (T425 lpxr, Chroma Technology), separating third and second harmonic generation or respective two-photon emission, thus allowing for simultaneous acquisition of two wavelengths with separate point detection on two photomultipliers for raster scan imaging. To detect THG signal, we collected the light through a 380–420 nm filter (ET400/40X, Chroma Technology, Vermont, USA), and to detect MBP immunostaining (secondary antibody Cy3), we used a 570–630 nm filter (ET600/60, Chroma Technology, Vermont, USA). Image stacks were acquired from 20 µm thick horizontal sections immunostained for MBP (1:200 rat anti-MBP, 1:200 goat anti-rat Cy3). Images were denoised using CANDLE(32) non-local means denoising software implemented in MATLAB (MathWorks), which can be found at https://sites.google.com/site/pierrickcoupe/softwares/denoising-for-medical-imaging/multiphoton-filtering.

### mRNA Synthesis for Blastomere Injections

To synthesize the mRNA for blastomere injections, GCaMP6s (Addgene plasmid 40753) and mCherry (plasmid gift of Dr Keith Murai) were each cloned into the pCS2+ vector. The GCaMP6s plasmid was cut with NotI / Klenow fill in / BglII, the mCherry plasmid was cut with BamHI / EcoRV, and the pCS2+ vector was cut with BamH1 / SnaB1. The plasmids were linearized with NotI, and the capped mRNA of GCaMP6s and mCherry were transcribed with the SP6 mMessage mMachine Kit (Ambion, Thermo Fisher).

### Calcium Imaging by Two-Photon Microscopy

Bilateral mosaic tadpoles expressing GCaMP6s at stage 48 were immobilized in 2 mM pancuronium dibromide (Cedarlane/Tocris 0693) and embedded in 1% low melting point agarose in a petri dish filled with 0.1X MBSH. For visual stimulation, 10 ms full-field light flashes at different frequencies were generated with a red luxeon LED placed next to the petri dish and controlled by an Arduino UnoR3 board (RRID:SCR_017284). Animals were imaged with a 20X water-immersion objective (1.0 NA) mounted on a commercial high-speed resonance scanner-based multiphoton microscope (Thorlabs) with piezoelectric objective focusing (Physik Instruments). GCaMP6s fluorescence was excited using Spectra Physics InSight X2 femtosecond pulsed laser tuned to 910 nm. Emission signal was detected with a GaAsP photomultiplier tube behind a 525/50 nm bandpass filter. Calcium signal was recorded from a single optical section (250 µm x 250 µm imaging field), focusing on the neuropil region of one tectal hemisphere. 20 s after initiating capture, the animal was shown a 1 min test stimulus (5 flashes of 20 ms duration, presented 15 s apart), followed by 10 minutes of continuous LED flashes at the chosen frequency of stroboscopic illumination (or no flashes for the spontaneous activity condition), immediately followed by another 1 min of the test stimulus. This was followed by a 5 min rest period between trials, then the stimulus sequence was repeated with the next strobe frequency being tested. The strobe frequencies tested were 0.0625 Hz, 0.25 Hz, 0.5 Hz, 1 Hz, 2 Hz, and 4 Hz.

### Quantification and Statistical Analysis

#### Calcium Imaging Analysis

Calcium recordings were registered using NoRMCorre(33), and analyzed using custom Matlab scripts. For each animal, the calcium signal was averaged over the neuropil region, the ΔF / F trace was calculated using the 20th percentile as baseline, then the signal corresponding to the strobe period was detrended with a fourth-degree polynomial before performing Fourier spectral analyses. The code for analysis of calcium responses to stroboscopic visual stimulation can be found at: https://github.com/RuthazerLab/Myelination_Strobe-Ca2-Analysis.

#### MBP-masked 3A10 Volume Analysis

Following immunohistochemical processing of tadpoles reared in strobe or ambient light for 7 d, image stacks of each section containing the optic chiasm were exported to Imaris 9.2 (Bitplane) for 3D reconstruction of the chiasm and analysis in a blinded fashion. First, sequential stacks for each optic chiasm per animal were imported into the same file and manually aligned for 3D visualization. For each stack, the surface creation function in Imaris was used to automatically construct a surface from the MBP channel, setting the background subtraction with surface detail at 0.118 μm and largest sphere diameter at 20 μm, thresholding automatically, with the final seed points approved by visual inspection. To study ensheathed axons at the chiasm, the 3A10 channel was masked with the MBP surface, and this masked channel was used to construct the MBP-masked 3A10 surface, using the same rendering parameters as the initial MBP surface. The volumes of the MBP-masked 3A10 surface were summed to find the total volume of MBP-associated axons at the optic chiasm.

#### Statistical analysis

Statistical analysis was performed in GraphPad Prism 8.0, and the details for each experiment can be found in the figure legend and figures. For the MBP-masked 3A10 volume, the ROUT test indicated one outlier (from 1Hz strobe group) that was removed.

